# AOPWIKI-EXPLORER: An Interactive Graph-based Query Engine leveraging Large Language Models

**DOI:** 10.1101/2023.11.21.568076

**Authors:** Saurav Kumar, Deepika Deepika, Luke T Slater, Vikas Kumar

## Abstract

**Background:** Adverse Outcome Pathways (AOPs) provide a basis for non-animal testing, by outlining and encoding the cascade of molecular and cellular events initiated upon stressor exposure, leading to adverse effects. AOPs are therefore considered an integral part of the New Approach Methodology (NAMs) to human health risk assessment. Over the last few years, the scientific community has shown immense interest in developing AOPs with crowdsourcing, with the results of those efforts being archived in the AOP wiki: a centralized repository coordinated by the OECD, which hosts nearly 512 AOPs. However, the AOP-wiki platform currently lacks a versatile querying system, which would enable developers to integrate and explore the AOP network data in a manner that would permit their practical use in risk assessment.

**Implementation:** This work proposes to expose the full potential of the AOP wiki archive by adapting its data into a Labeled Property Graph (LPG) database. In addition, the solution provides both database-specific query and natural language query interfaces to support retrieval and integration of the graph data, including the ability to chain stepwise queries. To evaluate the platform, a case study is presented, with three levels of use case scenarios (simple, moderate, and complex query). This tool is freely available on GitHub (InSilicoVida/AOPWiki-explorer) for wider community usability and further enhancement.

**Conclusions:** The presented query engine increases the potential of AOP exploration, by reducing the time, human resources, and technical ability needed to make practical use of the data they contain. Also, its dynamic prompt generation capability, will lead to precise query generation as the interaction increases. We, therefore, anticipate that the approach will be integral to the future realisation of new approach methodologies in chemical risk assessment.

## Introduction

The concept of Adverse Outcome Pathway (AOP) was defined in 2010 by Ankely et al. to streamline the idea of Next Generation Risk Assessment (NGRA)^1^. Ankely et al. described AOPs as an organizing framework to facilitate the representation of cascade events initiated by the stressor’s perturbation at the molecular level. Events in AOPs are broadly categorized into three sub-categories i.e., Molecular Initiating Event (MIE), Key Event (KE), and Adverse Outcome (AO), with the respective biological organization level based on their occurrence to effectively apply for risk assessment^1,2^. AOPs broadly capture the mechanistic knowledge of events in a well-connected sequential format, which helps in the strategic planning and development of new approach methodologies (NAMs) such as in-vitro tests, targeted assays, and Integrated Approaches to Testing and Assessment (IATA)^1–3^.

AOP developments are backed by guidelines and principles mentioned in the AOP Developers’ handbook, which are revised regularly to reflect feasible principles for improving the AOPs^4^. As per the mentioned guidelines, AOPs developed over the years are stored in AOP Knowledgebase (AOP-KB). AOP-KB stores machine-readable textual information of AOPs in XML (Extensible Markup Language) formatted data. On top of AOP-KB, there are two other services i.e., AOP Portal and AOP wiki, that act as query interface. The AOP portal enables the search of AOP and key events with keywords and also provides information about AOP endorsement. AOP-wiki is hosted as a central repository for AOPs developed as part of the OECD AOP development Programme. AOP-Wiki provides a platform to crowd-source and organize available knowledge as well as provides read/write access to AOP-KB following the OECD EAGMST guidelines. There are various third-party tools developed to enrich and support the AOP development which are assembled in the AOP-wiki and AOP-KB platform.

The ubiquity of graphs in data representation cannot be overstated. Any form of data can be reimagined as graph data whether it is tabular, relational, or even unstructured text^5^. In mathematical terms, graphs are represented with notation G = (V, E), where V is the set of nodes and E is the set of edges, where each edge contains a pair of nodes (u, v) representing a connection between node u and node v^5^. The implicit data structure of AOPs is a graph as it represents the cascade of events in the form of nodes (KE) connected by edges (KER). Building upon this implicit graph structure, initial efforts have been invested in modeling AOPs using RDF (Resource Description Framework) triples. RDF is one of the approaches for representing and modeling data in graph format. Notable examples include the work done by Ives et al. on systematically breaking down the key event term of AOP into three sub-terms i.e., process, object, and action. Each sub-terms were linked with respective ontologies and assigned a unique identifier^6^. Similarly, Martens et al. converted AOP data into Resource Description Format (RDF) by linking terms such as AOP, key event, and key event relationship with persistent and resolvable identifiers, which allows interoperability with other resources^7^. The introduction of RDF presents a paradigm shift in data modelling by encapsulating data as subject-predict-object triples, enabling a structured and semantically rich framework for describing AOP^3,7,8^. The adoption of RDF aligns with the principles of Data FAIRification, which advocates for data that is Findable, Interoperable and Reusable ^9^. RDF-modelled data is queried using SPARQL (SPARQL Protocol and RDF Query Language), which allows searching, filtering, and extracting of the desired information. RDF conversion makes AOPs querying possible up to a certain extent, but the complex and lengthy query of SPARQL makes it hard for the non-technical user to implement it. Also, querying the resources in SPARQL required the exact identifier (Uniform Resource Identifiers) of the nodes, which is troublesome to remember. The response to SPARQL queries is in tabular format in the AOP-WIKI RDF tool which is counterintuitive as per the graph nature of AOPs^3,7^ The tabular response gives a hard time for the user to understand, how various AOP elements are interconnected. It does not effectively illustrate how individual biological processes are orchestrated by multiple events within the complex network structure. Furthermore, SPARQL lacks the flexibility for deep-data traversals, and path analysis and has a high logarithmic cost of graph transversal^10–12^.

In this article, we unveil the capacity of a Labeled Property Graph (LPG) data modelling paradigm to serve as a natural data structure for Adverse Outcome Pathways (AOP). In LPG, data is organized into nodes and relationships in contrast with RDF-triples which consist of subject-predicate-object. We demonstrate how an LPG-adapted graph database empowers the risk assessors, and modelers to easily capture the multifaceted relationship that underlies AOPs, while efficiently exploiting the wealth of unstructured information associated with each AOPs. Crafting database-specific queries is a challenge, particularly for non-technical users. To alleviate this complexity, a pioneering step has been taken to integrate a powerful Large Language Model (LLM). The LLM model with its ability to learn textual patterns with few short prompts, bridges the technical gap and provides a user-friendly communication to extract value insights from graph data. Beyond simplifying query composition, the interpretation of extracted data in an interactive network further eases AOPs network in-depth analysis. To the best of our knowledge, none of the tools till now has implemented AOP data in its native format i.e., graph. This work provides a unified full-stack solution of graph data implementation that encompasses essential components i.e., data structure, query generator, and interactive interpretation for AOPs. This tool harmoniously converges to create an invaluable toolset for the AOP developer’s community.

## Materials and Methods

The method of adapting AOP wiki data in a graph structure, with multiple query interfaces, is broadly divided into 3 steps i.e., 1.) XML to graph conversion, 2.) Natural language to graph query and 3.) network visualization. All three components seamlessly interconnect to build a cohesive tool. In this interconnection the schema of the generated graph database in step 1 is useful for prompt creation, followed by query generation using the same prompt (step 2), and finally visual rendering of the network using query (step 3). In the subsequent methods section, we will delve deep into these 3 steps. Additionally, we will also discuss the container-based deployment of the tool.

### XML to native graph database conversion

XML version of the latest AOP data released in April 2023 was downloaded from https://aopwiki.org/downloads. The AOP data gets regularly updated (quarterly in a year). The downloaded XML file was then parsed into a Python dictionary object using the xmltodict library^13^. The converted dictionary object consists of elements, such as AOP, key-event, key-event-relationship, chemical, stressor, and taxonomy which are the building blocks of the whole AOP network (Fig 2). These data elements hold unique IDs, which helps in referencing and restructuring the block to develop the whole AOP network from scratch. Neo4j platform was used to implement LPG schema of AOP network^14^.

**Figure 1:**
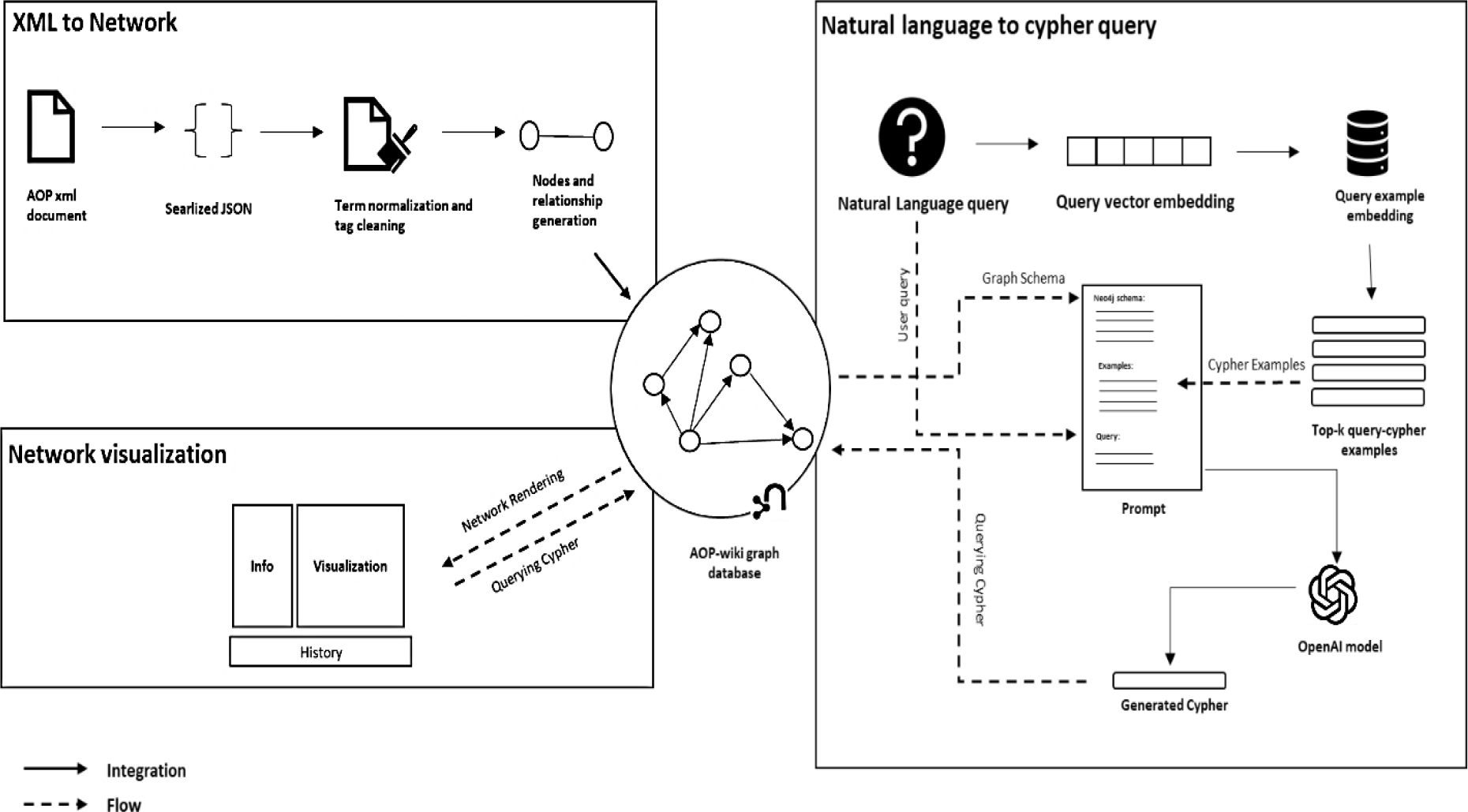
Workflow architecture of AOP-Wiki explorer. Each component mentioned in the workflow is developed separately but synchronizes with each other in the operational pipeline. XML to network conversion component processes and compiles the data in into a network form. The schema of compiled AOP-network database is assembled with the prompt to enable natural language to cypher generation process. Finally, the network visualization component retrieves the graph from the query and renders it in network form. AOP-wiki graph database acts as a central hub facilitating interaction among these components.

**Figure 1a:**
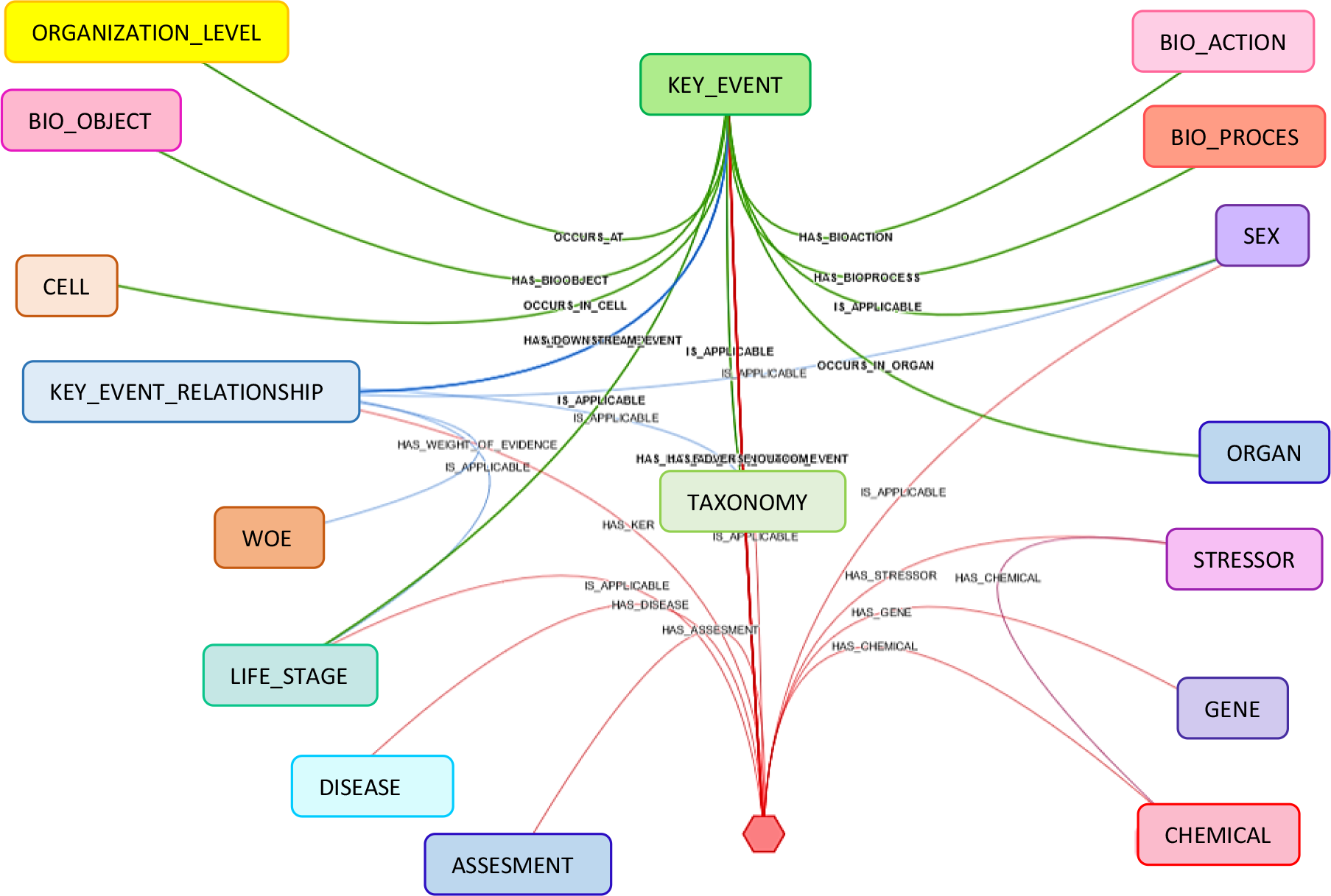
Neo4j graph data model schema for AOP-wiki XML data. Apart from key elements of AOPs i.e., key event, key event relationship and Adverse outcome other implicit information of AOPs that buried under rich-textual is extracted and explicitly represented as separate node.

**Figure 2:**
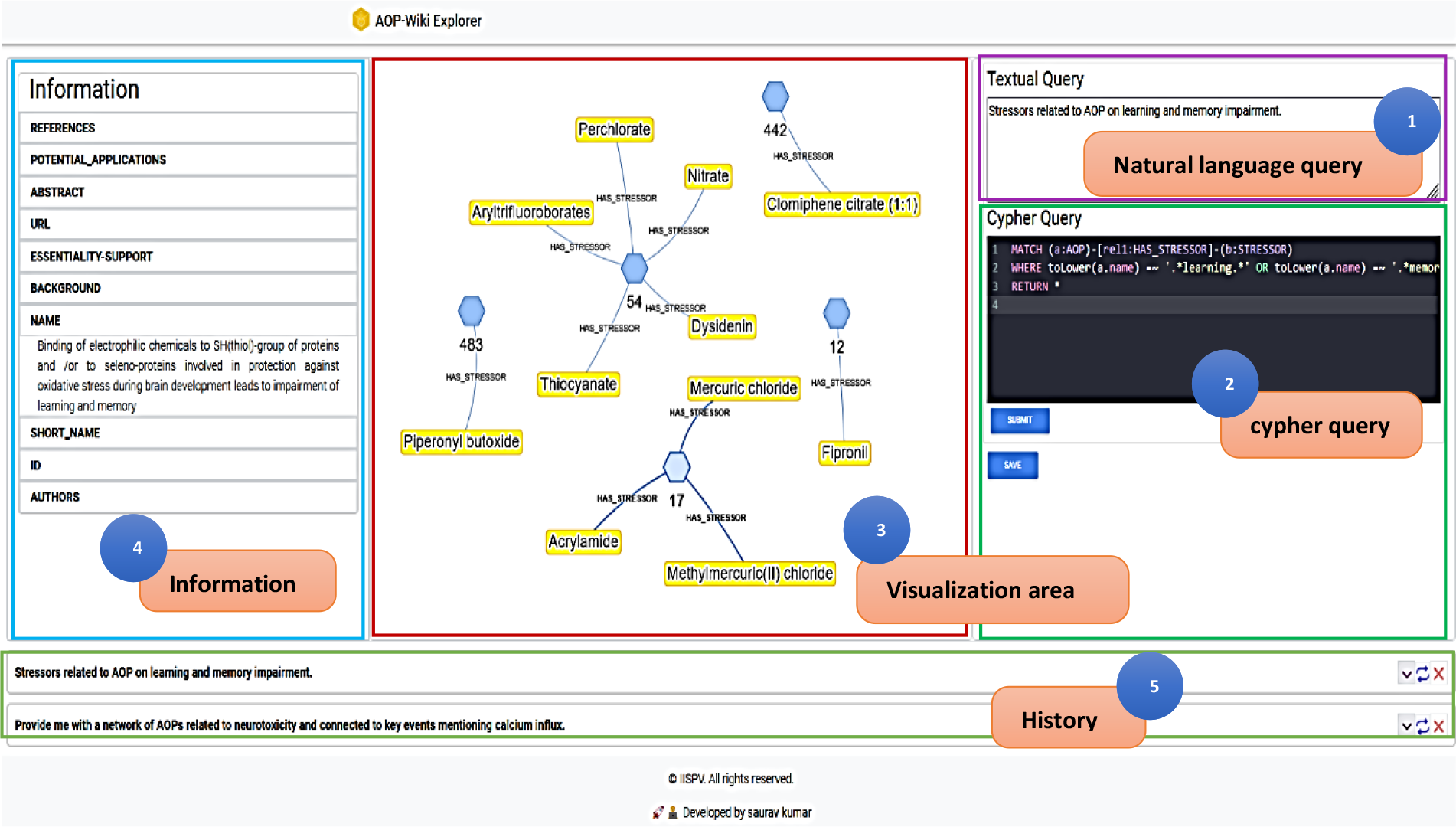
AOP-Wiki explorer user interface. The AOP Wiki explorer user interface provides an interactive playground to get graphical insights into AOP. The user interface rationally designs into 4 components i.e., 1) Natural language query area, 2) cypher query area 3) visualization area 4) information area and 5) history area.

Neo4j is a widely used graph database management system, designed to store, manage, and query large amounts of data organized in graph structure. Neo4j relies on cypher query language to interact with graph data. It is designed to express simple to complex queries, that can transverse and retrieve the relationship within the graph. Neo4j at its core, uses Java to run the engine, to make it accessible programmatically Py2neo library was used^15^. Py2neo is a Python library, which acts as an interface between Python and Neo4j and makes programmatic access to neo4j with ease. With Py2neo, nodes, and relationships are built while keeping a consistent naming convention of node and edge labels and their properties. Nodes and edge labels are in uppercase and the label having more than one word are joined with the snake naming convention. Properties of nodes and edges are written in lowercase. The strict naming convention provides robustness and consistency in further integration.

The graph schema in Figure 2 illustrates the way information within AOP-wiki is structured and represented as a graph data model. Typically, in AOP graphical representation, there are mainly three types of nodes i.e., AOP, key event, and Key event relationship. In this schema, it has been extended to show information like taxonomy, sex, biological organization level, assessment methodology, and many more. Also, while populating the properties of nodes, textual information was processed using regex to clean the HTML tags to make the text human readable. The extended schema gives flexibility to the user to retrieve information at different levels of granularity within AOP, facilitating in-depth analysis. Some of the crucial textual information of AOP such as their assessment methodology and weight of evidence are also represented as nodes to increase information accessibility. The terms like “life stages” and “sex” underwent processing to eliminate redundant information and were standardized by assigning them persistent identifiers from the referenced ontology. From the AOP wiki portal, sections containing descriptive textual information about each AOP, such as abstract, description, the weight of evidence, and assessment method methodology, etc. have been combined into a single document. Over the combined document Named Entity Recognition (NER) is applied to extract the biological concepts such as gene, protein, chemical, and disease using BERN2 biomedical named entity recognition and normalization tool^16^. Normalized entities are mapped to respective databases and integrated into the network as nodes.

### Natural Language to graph query

To bridge the technical gap and provide flexibility to query information in natural language form, an infamous OpenAI’s GPT-4^17^ model was used. OpenAI’s GPT-4 is a LLM designed to understand and generate human-like text. The GPT-4 model is trained on an extensive and varied corpus of text derived from the internet, encompassing sources like books, articles, websites, documentation, and code repositories. Training on such a massive amount of data enables the model to learn patterns, syntax, semantics, and contextual nuances. GPT-4 model inherently does not possess specific knowledge about cypher queries, instead, they acquire knowledge through extensive training data or prompts they are exposed to. Description and examples of cypher query provided in the prompt enable the model to generate coherent patterns. However, the response from the model is based on statistical patterns they have learned while training, rather than a deep understanding of the Neo4j cypher.

The quality and correctness of generated cypher queries depend on how precise the prompt is provided. The precise prompt here signifies explicit instruction, relevant context, illustrative examples of desired responses, and clear specification of the output format. To build precise prompts, in this work dynamic few shot prompt generation methodology has been adopted. In dynamic methodology, a few examples in the prompt are provided similar to the query asked by the user. Similar examples make the model aware of the syntax and variables to be used for generating the cypher. Similar examples were stored in the form of vector embedding generated by “all-MiniLM-L6-v2” sentence transformer model^18^. Chroma vector database^19^ was used to store examples of embedding. This database gets updated with more examples as queries are saved by the user. The variable part in the dynamic prompt is the provided example query and query itself, whereas other components are static. Static components of prompts include graph schema, context, and output format. Graph schema holds information about how the database is structured and the properties mentioned in nodes and edges. Neo4j module of langchain^20^ was used to generate the schema of the graph database. Context provides precise instructions to the model, while output format provides schemas such as JSON, or YAML which enables to generate machine parseable cypher.

The overall compilation of prompts follows the sequence mentioned in Figure 1. First, the real-time query of the user is embedded. Based on this query embedding, similar example queries are retrieved using cosine similarity from the vector embedding database. The extracted examples are then merged with other components of the prompt and fed to the model for the final cypher query generation.

### Interactive network visualization

For network visualization, an interactive user interface was built called AOP-wiki Explorer (Fig 3). The interface is thoughtfully structured into five distinct components: i) natural language query, ii) cypher query, iii) visualization area, iv) information and v) history. These components orchestrate harmoniously and provide users with an adaptable playground to analyze AOPs. Various components and their role are briefly described as follows:

#### Natural language query

It takes user-initiated queries in natural language form. Input to this component is mandatory as it helps to keep track of human-understandable queries. Additionally, it assists in generating cyphers for users who may not have technical knowledge about cypher queries.

#### Cypher query

Takes direct Neo4j cypher query as input. Here users with technical knowledge can craft intricate cypher queries and are able to leverage the full potential of a graph database to extract precise insights and patterns. Syntax highlighting is supported for the cypher that provides readability to the code.

#### Visualization Area

It is a visual output component that renders the pattern given by the cypher query. Visual representation empowers the users to unravel complex relationships, dependencies, and trends within graph databases (in this case AOPs). Visualization is powered by the Neovis library^21^, which is part of the Vis JavaScript library specialized in handling Neo4j data.

#### Information Area

It provides comprehensive details stored as properties of nodes and edges. When tapped over the network component (nodes and edges) in the interactive visualization, it shows the tabular list of properties with summarized information (in this case summary information extracted from AOP-wiki). It also provides the URL to navigate the source of information i.e., AOP-wiki.

#### History Area

This component allows users to retrace their steps and revisit previous queries. This feature facilitates quick referencing, iterative exploration, dynamic data investigation and query chaining.

### Container Based Deployment

Orchestrated container-based deployment has been adopted in this work. Four distinct containers i.e., Jupyter, frontend, backend and neo4j operate synchronously under the same network. With a single command i.e., “docker-compose up”, the entire system will be up and running without any external dependencies, offering a jupyter interface to populate the database and frontend access to query it. It effectively abstracts the complexity of deployment, making it easy to expand the database with more information. Containerization provides a seamless way to update the tool with the quarterly release of AOP-wiki XML data.

In this work processing of raw data to the network has been done with Python3^22^. The web app was developed with React^23^, a java script framework for UI development. Along with this other java script library such as Neovis.js, codeMirror.js, etc. were used to support other components for visualization. The backend was developed with a flask micro server in Python. Each component was constructed as a Docker container^24^, and their coordination is managed through Docker compose version 3.7.

## Result and Discussion

### Adopting AOP-wiki XML as a Graph database

This work adapts AOP-Wiki XML information into an LPG data structure. Transforming XML data to graphs enables the researchers to query AOPs in their more natural way. Along with this AOP wiki information has been further enriched with information about genes/proteins, chemicals, and diseases mentioned in each AOP using NER. A total of unique 1135 biological entities have been identified, and out of that 438 were gene/protein, 377 chemicals, and 320 diseases (Supplementary file 1). Each of the entities was attached to their respective ontology such as CHEBI^25^ is associated with chemicals, MIM^26^ is related to disease, NCBIGene^27^ for genes, and MESH^28^ is a general purpose which covers multiple domains. With ontology association, the problem of duplication of the same entity with a synonym has been resolved. The enrichment allows users to search AOPs with the very fundamental information of their work such as genes or chemicals associated with them and gives more flexibility to information retrieval. To make the AOP graph data free of redundant information and FAIRER, information such as life stages and sex has been filtered and attached to its consistent identifier. Out of the 23 identified life stages, redundant ones like “Adults” and “Adult,” as well as “Foetal” and “Fetal,” have been filtered and attached to their permanent identifier. Similarly, lots of other terms like “Adult, reproductively mature”, “1 to < 3 months” and many more were carefully assigned with specific ontology and definition. Terms lacking specified ontology remain unchanged. FAIRified life stages are available in supplementary file 2. In addition, with FAIR data, it is equally vital to have a well-defined data model to facilitate the accessibility of information at a detailed level.

The schema of the graph data model was crafted to align with criteria aimed at enhancing information accessibility and interoperability, thus leading to broader applicability of the tool. Information such as life stages, sex, taxonomy, gene/protein chemical, etc. of AOPs and KE often reside deep in their textual descriptions, so rendering and filtering information on these criteria is challenging. Representing this information as separate nodes leads to fine-level granularity, reduced term redundancy, and wider accessibility of information. Additionally, representing KERs as edge property between two KE limits the accessibility of other vital details of KER such as sex, life stages, etc. To address this concern, multiple schemas were designed and the one which meets the criteria was selected as shown in Figure 1, the network visuals of all other tested schemas are present in supplementary file 3 (Figure S2 and S3).

So far, our conversation has been centered around the process of transitioning to a native graph database to adapt the LPG schema. Going forward, our discussion will primarily shift its focus towards the reasons for proceeding with this transition, highlighting its advantages with some examples and use-case with respect to AOPs, while also pointing out the limitations of the current implementation in a future perspective. To begin, let us provide examples of well-known biological databases that adopted this transition, such as Reactome^29^, GREG^30^, GeNNet^31^, HRGRN^32^ etc. These databases have made this shift in response to the demand posed by biological complexity and extensive interconnections between different biological resources. Storing biomedical information in a relational paradigm (SQL/NON-SQL) has been widely practised due to its technological maturity in the database field^10^. However, data such as AOPs, protein-protein interaction, disease-chemical interaction, and many others like that hold densely connected relations. Querying densely connected information requires resource-intensive JOIN operation in relational databases which impact the performance as the data grows^10^. Although the tabular data can be modelled in graphs but optimizing complex queries and graph algorithms like path traversals are challenging without complementary adjacency list^10^. Graph database system such as Neo4j offers direct interaction with native graph model, eliminating manual relationship engineering^10,14^. They provide straightforward methods to access, curate and operate on data while also offering flexibility to adapt and evolve with the introduction of new data or schema. Furthermore, it is essential to acknowledge the significance of network analysis in unravelling the intricacies of biological representations. Villeneuve et.al have published a paper demonstrating, how network analysis can be effectively utilized in AOP development^2,33^. Their work demonstrated the implementation of graph topology to declutter and refine the AOP network subgraph, and the application of algorithms such as betweenness centrality, topological sorting, and many more in the context of AOP networks. We recommend this article for comprehensive insights into the application of network analysis for AOPs.

### Addressing query crafting with LLM integration

Each database comes up with its syntax or query language tailored to optimize information retrieval. Declarative query languages, such as SPARQL and cypher, abstract implementation details enabling users to specify what data they want to retrieve. In contrast, imperative query languages require users to specify both what data to retrieve and how to retrieve it. SPARQL is employed for information structured in RDF (Resource Description Format) schema, while Neo4j graph databases utilize the cypher query language. Both cypher and SPARQL share their origins with structured query language (SQL)^34^. However, cypher chooses to represent query patterns in the form of a graph using ASCII-art syntax, whereas SPARQL follows a text-rich query format. The adaptation of the right query pattern reflects a lot of differences in the usability and understanding of the query. To illustrate the difference, let’s use the syntax provided in Figure 5 to retrieve information about chemicals associated with “fibrosis” as an adverse outcome using both cypher and SPARQL queries. Cypher accomplishes this query in just 5 lines of instructions, while SPARQL takes 16 lines, nearly three times to retrieve the same result. In addition, SPARQL also needs consistent node identifiers such as “dc: identifier”,” dc: title”, and “nci: C54571” to query the same information. The choice of the identifier depends on the ontology and schema adopted for data modeling.

**Figure 4:**
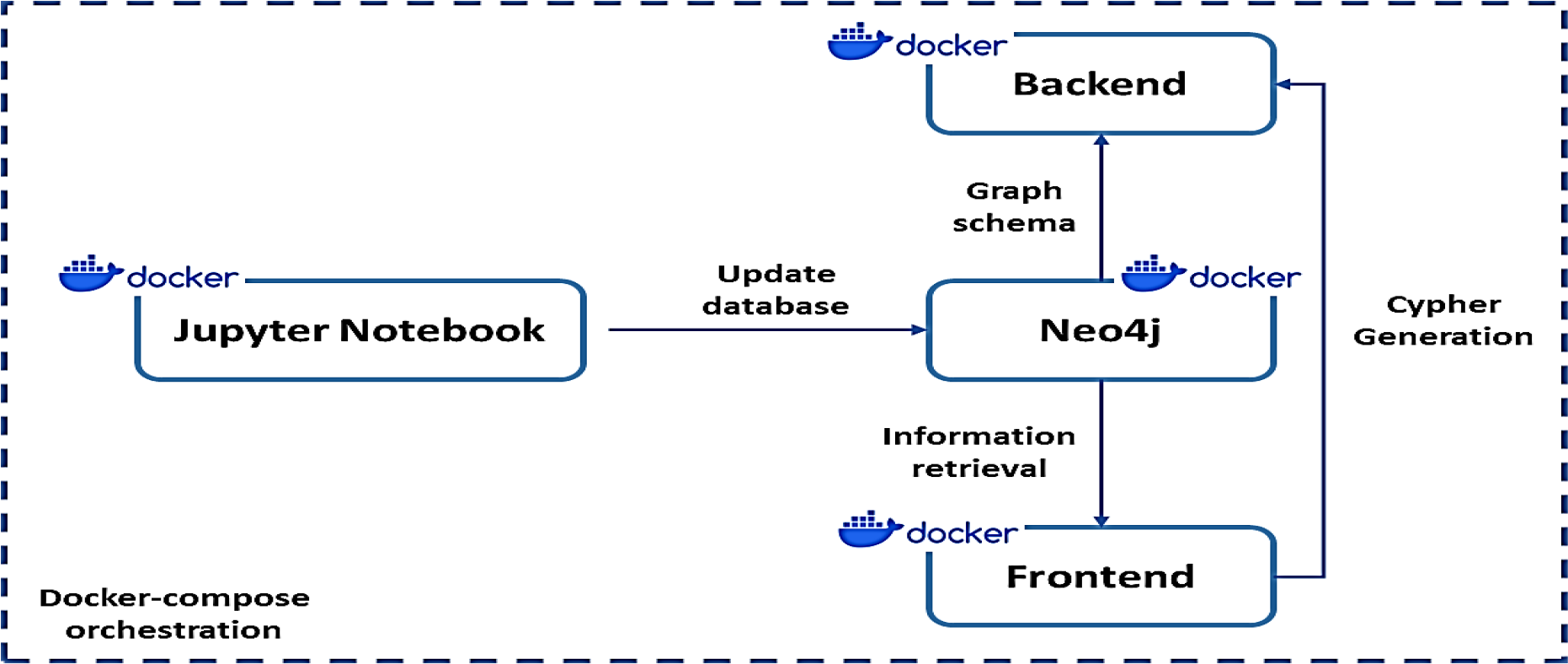
AOPWiki explorer deployment is orchestrated using container-based technology. A jupyter notebook will be accessible to users via their web browsers, providing them with the ability to add, edit, or remove information in the Neo4j database. The front-end component will enable users to query information, offering the choice between using cypher queries or natural language. The backend component will operate a micro-server tasked with translating natural language queries into Cypher queries, utilizing database schema and the user’s query.

**Figure 5:**
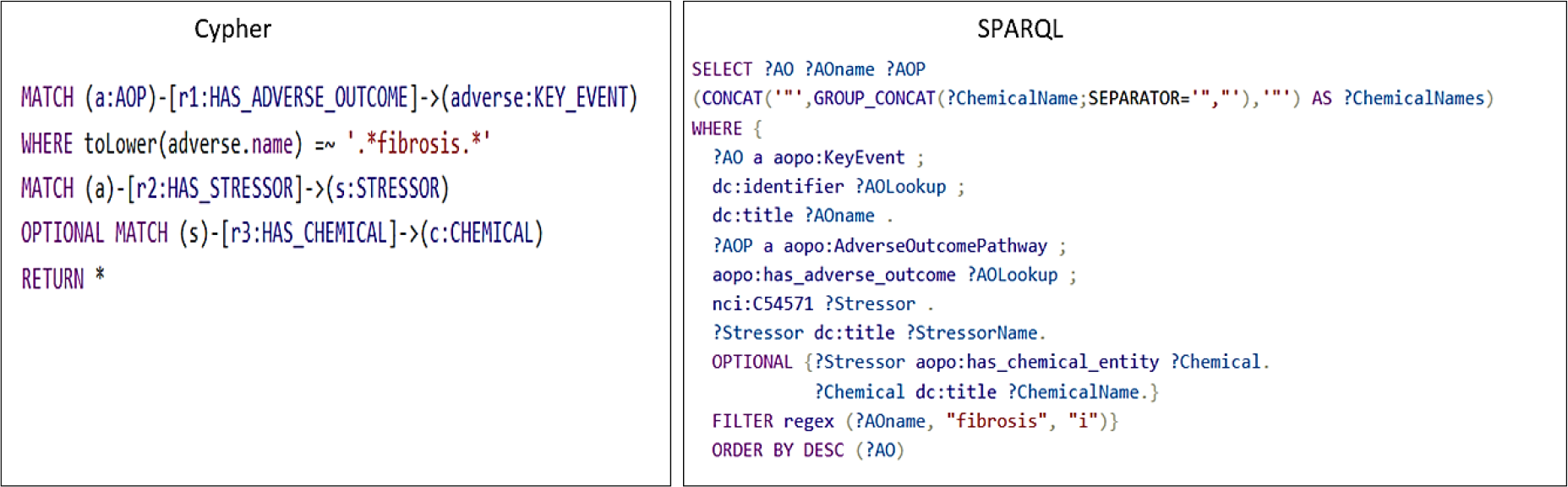
Query complexity comparison between cypher and SPARQL. Cypher and SPARQL queries are tasked to retrieve the chemicals from AOPs, which can lead to fibrosis. AOP-wiki RDF adaptation was used to query information via SPARQL.

Crafting complex queries either using cypher or SPARQL is not an easy task for researchers from a non-technical background. To bridge this technical gap, LLM comes in handy. With a few examples of natural language queries and cyphers, the model adapts the pattern and syntax of the cypher related to the database schema. This enables the model to generate cypher queries in real-time in response to the user’s natural language queries. A well-curated natural language query in accordance with the schema of a graph database is enough to generate a precise cypher. Having no communication barrier between the information source and the users will increase the accessibility and usability of the tool. The growing usability of the tool results in the addition of diverse queries in the AOP context from the users leading to an increase in precision and accuracy of cypher generation. The diverse queries asked by the developers will also reflect the needs and expectations they have from the AOP wiki; hence this will be also helpful in modulating the AOP wiki as per the user’s requirement in future updates. Table 1 illustrates the diversity of questions and queries researchers may need to address while building or evaluating the AOPs.

**Table 1:**
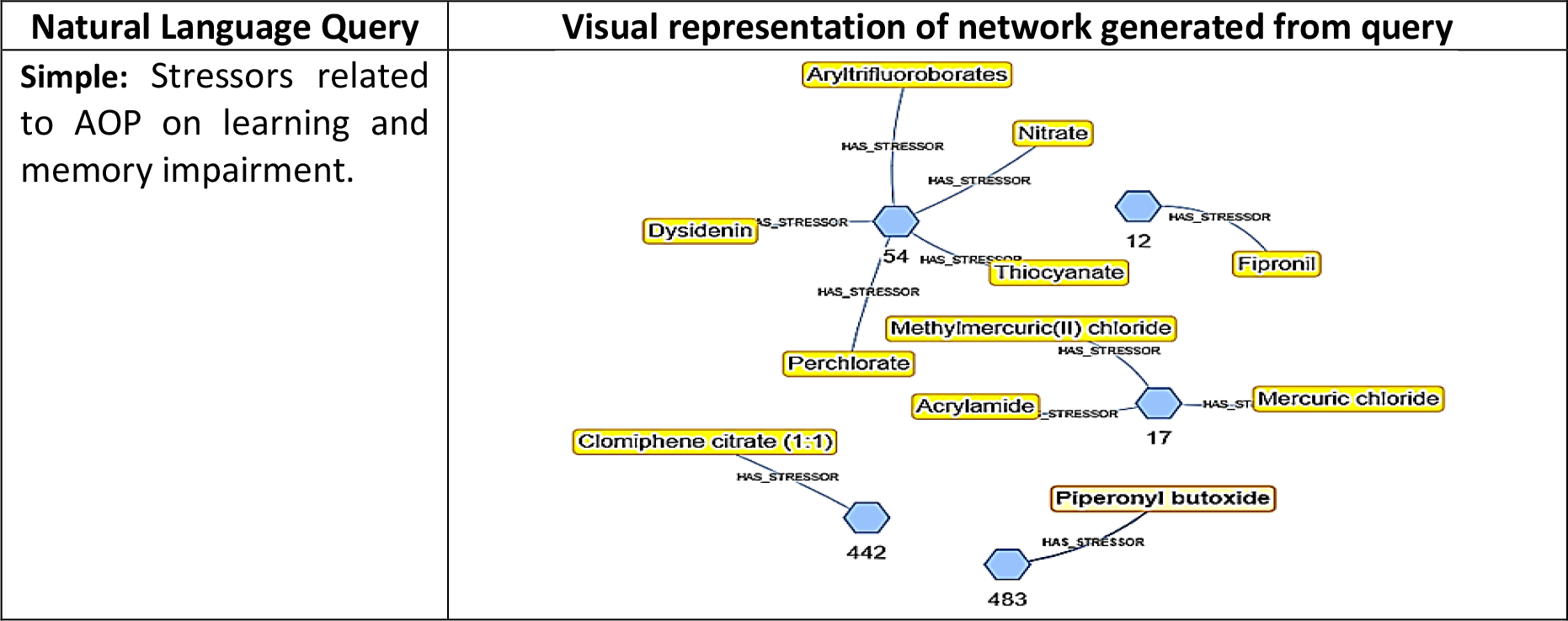
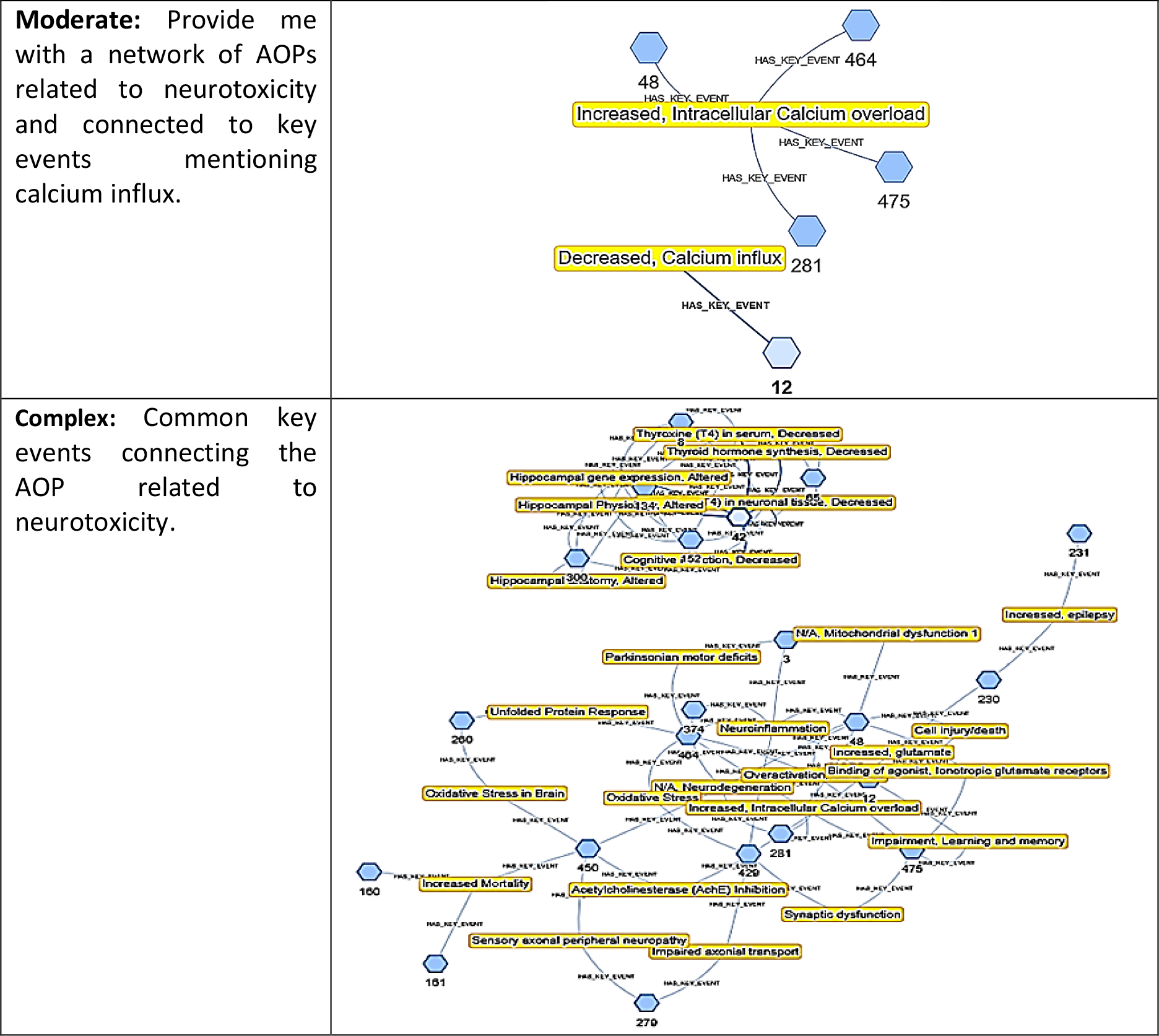
Different types of queries: simple, moderate, and complex are shown in natural language and network format. Cypher query for each natural language query is present in the supplementary 3 (Table S1).

Bridging the technical gap comes with the cost, in this study, we used OpenAI GPT-4 with 8k context size closed source LLMs, for each request to the language model charges $0.03/1k tokens. On average our prompt consists of 5k tokens, which costs us roughly $0.15 for each natural language to cypher generation. These expenses are anticipated to reduce over time, as the database accumulates relevant examples, we expect to rely on prompt examples resembling user queries, thereby reducing the need of imputing multiple examples. Furthermore, with the rapid advancement of LLM development in the open-source community, the problem concerning cost will be addressed soon.

### Interactive and stepwise (chained) query

The interface for exploring AOP development is designed to facilitate interactive and query chaining, acknowledging the inherent exploratory nature of AOP development. In this section, we will explain queries chaining with the help of an example query i.e. “What are the key events that are connecting the AOPs which are applicable to fish and rat taxonomy and have adverse outcomes related to fertility or reproductive issues”. We can infer that the complex query can be broken down into multiple sub-queries such as:

1. AOPs which are applicable to “fish” and “rat”.
2. Filter the AOPs, which mention “fertility” or “reproductive” issues in the adverse outcome.
3. common key events between filtered AOPs.

The above-mentioned queries are rationally divided into steps and queried separately, which allows users to analyze the results of the query individually and chain it with other queries if required. In Table 2 above mentioned natural language query is presented in a visual network and their cypher query is presented in supplementary file 3 (Figure S1). We can see that the network can be generated about different key events considering AOPs related to fertility or reproductive issues for fish and rats. Overall, the AOPwiki explorer can be helpful in extracting the information from AOPs with multiple queries enabling the user-friendly interface with the option to save past data.

**Table 2:**
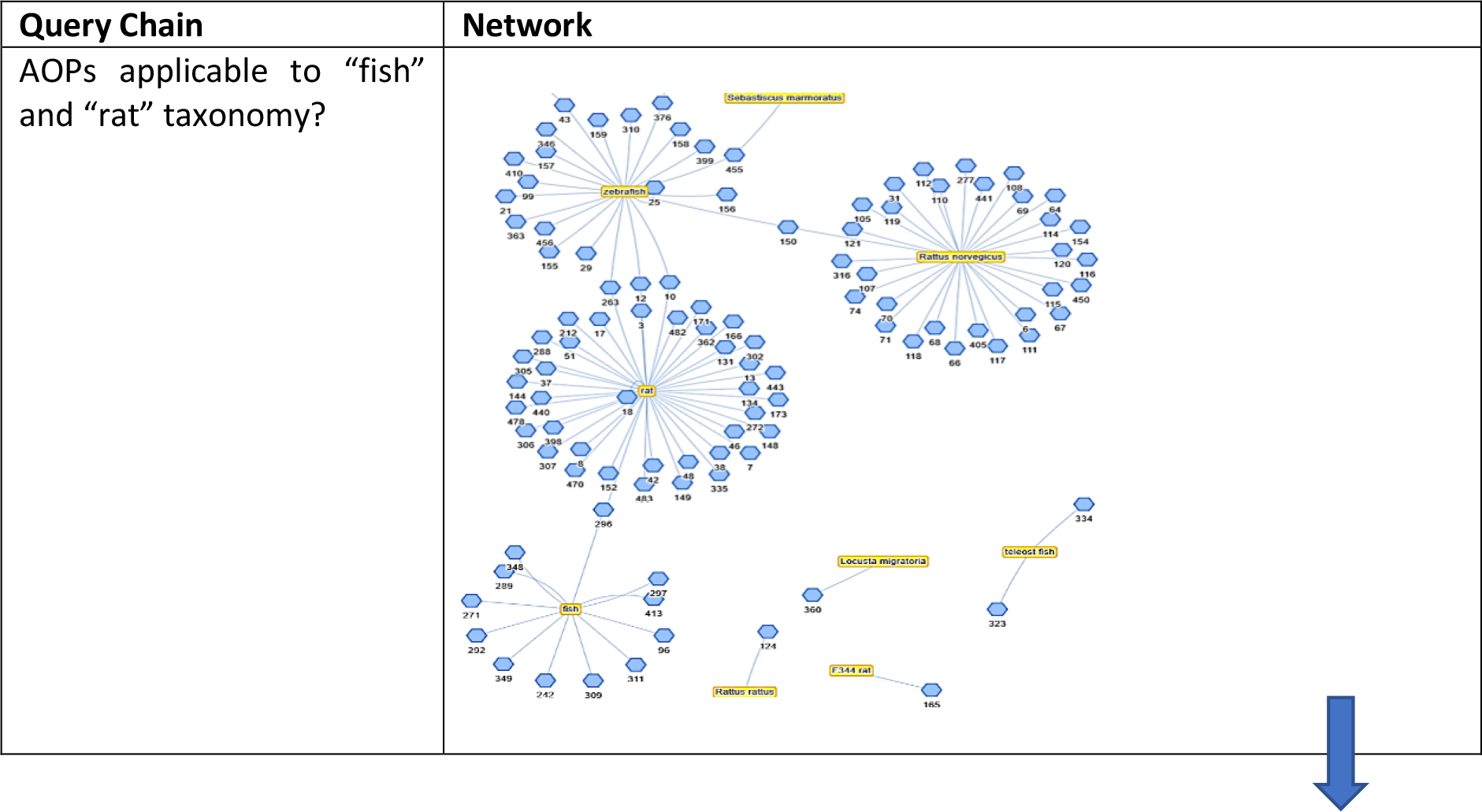
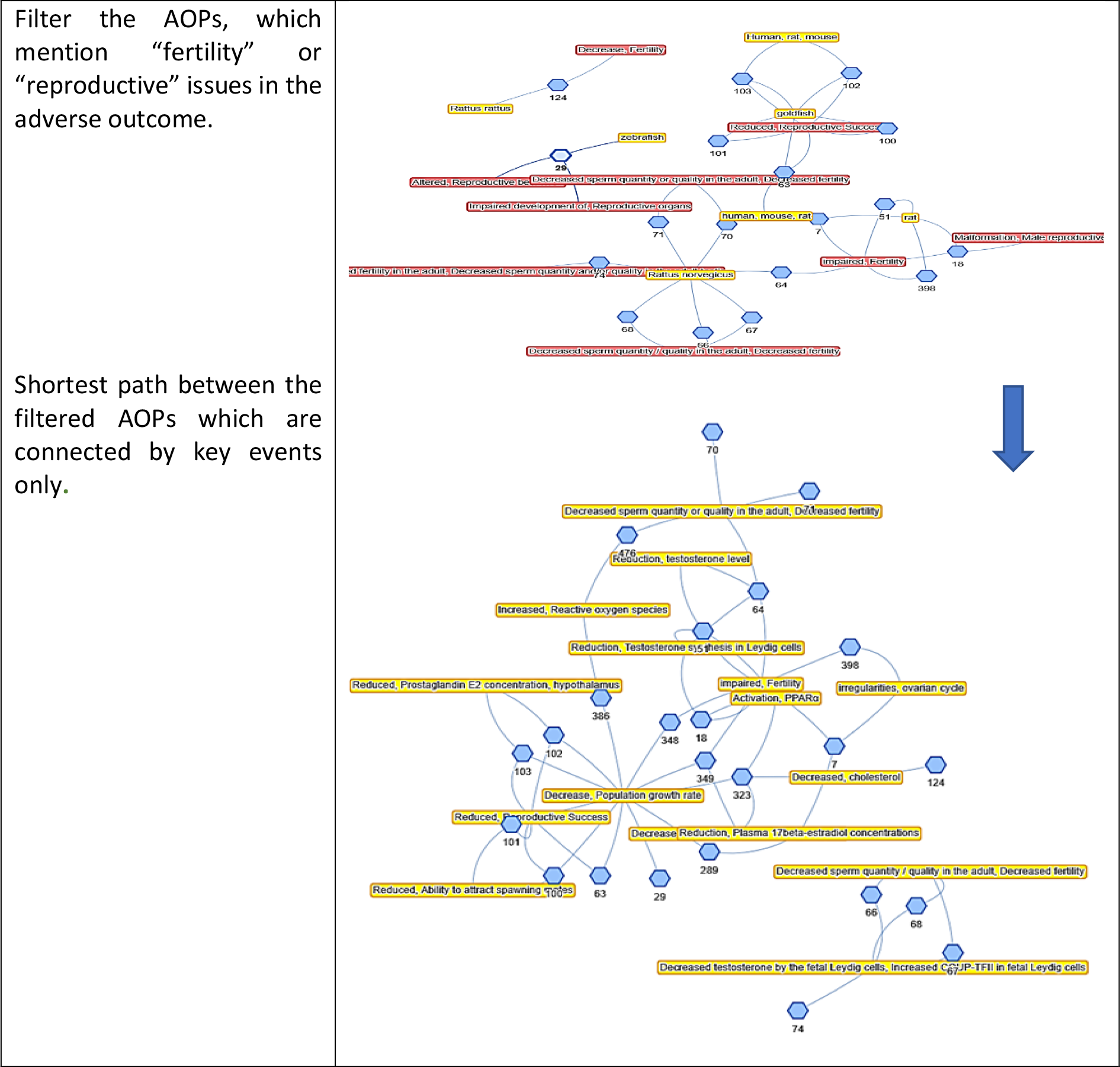
The table shows chains of queries. It shows how one query can be linked with another to pull out complex related information from the database. In this example, the initial query “AOPs applicable to “fish” and “rat” taxonomy?”, all AOPs having mentioned “fish” or “rat” in their applicable taxonomy can be pulled, chaining it with filter queries to select only those having “fertility or reproductive mentioned” in their adverse outcome. It was further narrowed down to extract the shortest path between the filtered AOPs, which is only connected by the key event.

### Future perspectives

The work presented herein provides an enhanced ability to computationally query, retrieve, and integrate AOP-Wiki’s data by converting it into a more amenable graph format, and by leveraging LLM to expose the internal representation of the AOPWiki database in a formulation more natural to AOPs’ formulation. However, the machine actionability of AOP-Wiki remains limited. This is due to the incomplete semantic definition and functional linking of its internal representation. While, as previously discussed, there have been efforts to transform AOP data into RDF, its natural formulation remains that of a relational database, with limited linkage between its internal entities and external semantic definitions.

AOPWiki and associated projects provide limited ontology-like structures, however these are not strictly defined, for example with species hierarchies not adhering to subsumption, meaning that e.g., information concerning mice may be split across distinct subsets of the ontology classified as “mouse,” “mouse and human,” “mouse and rat and human” (Figure 6). Outside the representation of its structured data, AOPWiki data are also highly unstructured, with much of the valuable and specific information being expressed in text.

**Figure 6:**
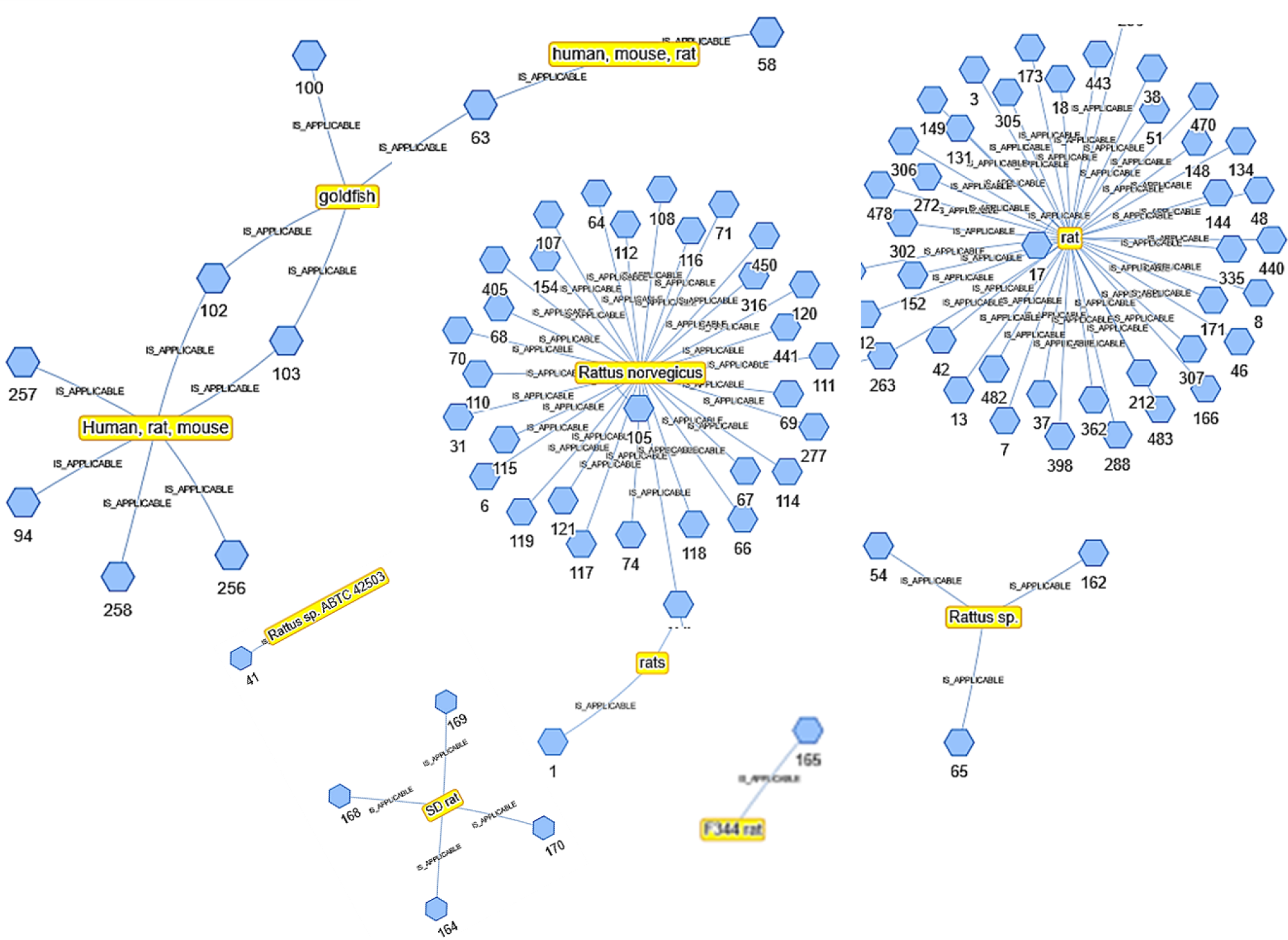
Redundancy of taxonomy in AOPs network, highlighting the absence of standardized approach in AOP wiki. The taxonomy of rats has split across distinct subsets such as “Rat norvegicus”,” rats”, “SD rat”. Also, it has been grouped with other taxonomy instead of single distinct node such as “Human, rat, mouse”. Additionally, inconsistencies occur in how the node name has been handled, as seen in cases like “Human, rat, mouse” and “human, mouse, rat”.

Importantly, the ability of LLMs to make sense of this representation is limited, and its performance in tasks involving identifying semantic entities from text or databases is limited^35^. Issues with hallucination, especially at the edges of knowledge are also well documented^35^. These issues are compounded where data are novel, on the edge of the scientific frontier, and therefore they cannot in this context be relied upon as an alternative for serious, collaborative efforts to producing high quality, FAIR, functionally linked data. We therefore see a more comprehensive approach to developing a fully semantic representation to be the next step in fully realising the potential of AOPs.

The value of building a fully semantic implementation of AOPWiki inheres in the ability to computationally integrate the knowledge expressed in AOPs with knowledge concerning the relevant entities across the scientific knowledge landscape. Increasingly, data concerning biomedical and chemical entities are made public in linked representations, most often using OWL ontologies^36^, which provide the ability for semantic confirmation of equivalence of entities across databases, to associate AOP events with external scientific evidence, and ontologies can be used to support the development of formal modelling approaches that will be required to use AOPs in risk assessment.

The process of realising this vision, however, will take substantial work. In this work, LPG implementation with its easier adaptability and graph analytics solution, tried to some solutions. However, in terms of semantics, RDF is a well versed and widely adopted by the community over the years, whereas semantics capability of LPG might lack. Some of recently developed LPG platform such as Neo4j provides a solution to integrate semantics in the LPG itself and provide the interface which can facilitate the enhanced user accessibility, query flexibility and scalability of AOPs database^37^.

Similarly, Scigraph^38^ is another tool, which supports the LPG backend for storing ontology-linked resources and is being used in large-scale projects such as Monarch initiative^39^, Human Brain Project^40^, SciCrunch^41^ etc. It will require innovation to align semantically enabled AOP’s information with other resources found elsewhere in the chemical and biomedical domains. Furthermore, while AOPWiki currently provides the ability to annotate its internal entities, such as events, with ontology terms, the nature of those relationships is not well defined, and is therefore not machine actionable. We envision a future where AOPs databases are designed to have a fully functional linked evidential association for AOPs. However, the data and metadata structures for these entities do not yet exist. While solutions such as BioPAX ^42^and BioLink^43^ would provide the features required to express associations, however, they will require additional work to support functional integration with AOP data structures, as well as the provenance, authorship, levels of confirmation, and other relevant metadata. These would also support efforts to perform information extraction, linkage, and integration from text and other unstructured or semi-structured data sources, both from within the AOPWiki database, as well as from other sources such as literature or policy documents. Knowledge representation like BioLink also supports the adaption in Labeled property Graph and with integration of NEO4J and semantics BioLink can provide semantically and algorithmically optimized solutions. Inspiration may also be drawn from other biomedical projects, such as the Gene Ontology Project^44^, whose instance databases include evidence codes. A general solution to evidence would support a wide range of uses across science, for example in the area of disease-phenotype associations and their use in biomedical research and healthcare.

## Conclusion

In this work, we represent AOP-wiki data as a graph and build a tool to enable the users to query and link information in a very flexible way. In this work, we filled the technical gap and implemented a solution using the LLM’s ability to generate cypher queries from simple natural language queries provided in the context of a graph database schema. To provide an exploratory platform to the AOP development process, we thoughtfully designed the interface which allows users to retrieve their complex research queries in multiple steps and keep track of individual queries separately. The results of the queries are retrieved in an interactive network form which can further be analysed and explored with the linked data to its nodes and edges. The detailed information on AOPs, key events, and their relation are also accessible in tabular form and provides the functionality to retrace the information source. This work also provides a prototype framework for the future development of AOPwiki or community databases.

## Supporting information

Sumplementary-3

BiologicalEntity -1

Life stages

## Acknowledgments

The work done here has been supported by the Partnership for the Assessment of Risk from Chemicals (PARC) funding by the European Union’s Horizon Europe research and innovation Programme under Grant Agreement No. 101057014.

## Code and software

Code is available through open source and MIT license at InSilicoVida/AOPWiki-explorer. Link to the tool (AOPwiki explorer) will also be added to the GIThub for public access.

