## Supplementary material for "AOPWIKI-EXPLORER: An Interactive Graph-based Query Engine leveraging Large Language Models": Sumplementary-3

**Table S1: Different variations of query i.e., simple, moderate, and complex are shown in natural language and cypher format.**

| Natural Language Query | Cypher Query |
|---|---|
| **Simple: Stressors related to AOP on learning and memory impairment.** | MATCH (a:AOP)-[rel1:HAS_STRESSOR]-(b:STRESSOR)<br>WHERE toLower(a.name) =~ '.*learning.*' OR toLower(a.name) =~ '.*memory.*'<br>RETURN * |
| **Moderate: Provide me with a network of AOPs related to neurotoxicity and connected to key events mentioning calcium influx.** | MATCH (a: AOP)-[rel1:HAS_KEY_EVENT]-(b:KEY_EVENT)<br>WHERE toLower(a.name) =~ '.*neuro.*'<br>WITH a<br>MATCH (a)-[rel2:HAS_KEY_EVENT]-(c:KEY_EVENT)<br>WHERE toLower(c.name) =~ '.*calcium.*'<br>RETURN * |
| **Complex: common key events connecting the AOP related to neurotoxicity** | MATCH (a:AOP)-[:HAS_ADVERSE_OUTCOME]-(c:KEY_EVENT)<br>WHERE toLower(a.name) =~ '.*neuro.*'<br>WITH collect(a.id) as nodes<br>MATCH (aop1:AOP) WHERE aop1.id IN nodes<br>MATCH (aop2:AOP) WHERE aop2.id IN nodes AND aop1<>aop2<br>MATCH path = shortestPath((aop1)-[:HAS_KEY_EVENT*..2]-(aop2))<br>RETURN path |

**Figure S1:** This example shows how one query can be hooked with another to pull out complex relation information from the database. In this example the initial query "AOPs applicable to "fish" and "rat" taxonomy?", all AOPs having mentioned "fish" or "rat" in their applicable taxonomy can be pulled, chaining it with filter queries to select only those having "fertility or reproductive mentioned" in their adverse outcome. It was further narrowed down to extract the shortest path between the filtered AOPs, which is only connected by the key event.

```
// AOPs applicable to fish and rat
MATCH (a:AOP)-[rel:IS_APPLICABLE]-(b:TAXONOMY)
WHERE toLower(b.name) =~ '.*fish.*' OR toLower(b.name) =~ '.*rat.*'

//Filtered AOP having fertility and reproductive issues as AdverseOutcome
MATCH (a)-[rel2:HAS_ADVERSE_OUTCOME]-(c:KEY_EVENT)
WHERE toLower(c.name) =~ '.*fertility.*' OR toLower(c.name) =~ '.*reproductive.*'
WITH collect(a.id) as nodes

// Match these AOPs together with key event as intermediate nodes
MATCH (aop1:AOP) WHERE aop1.id IN nodes
MATCH (aop2:AOP) WHERE aop2.id IN nodes AND aop1<>aop2
MATCH path = shortestPath((aop1)-[:HAS_KEY_EVENT*]-(aop2))
RETURN path
```

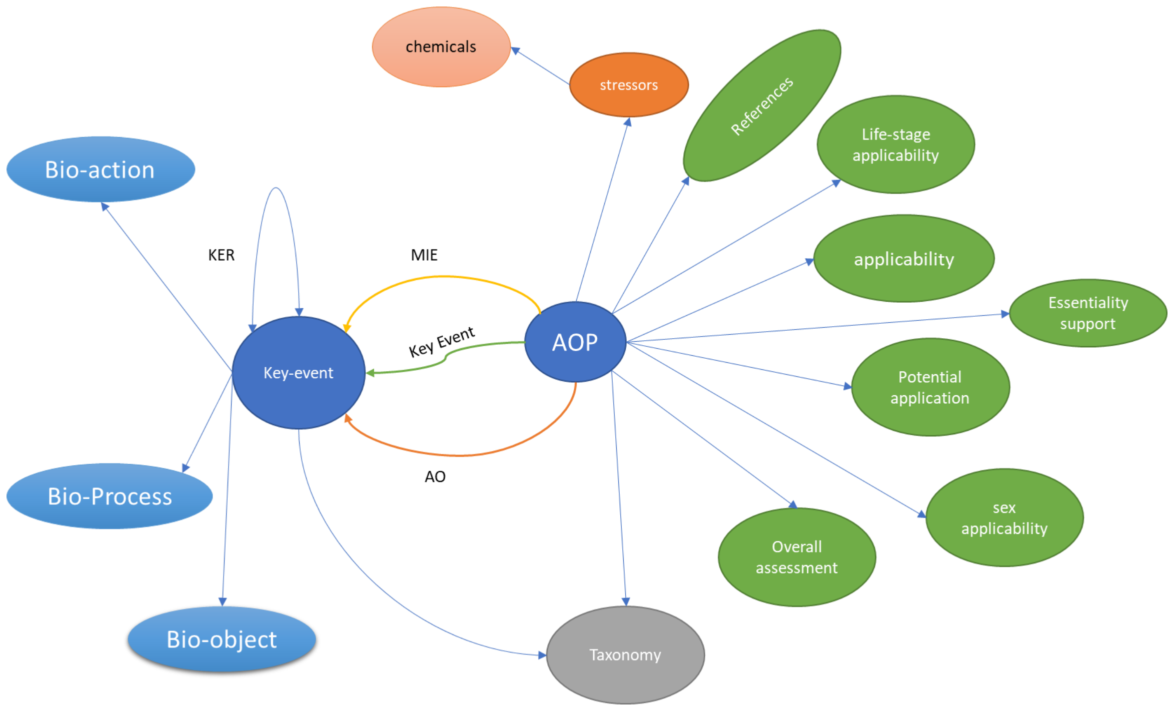


**Figure S1:** First tested schema of AOP, In this schema, Key event relationship information is stored as edge property of key-event edge.

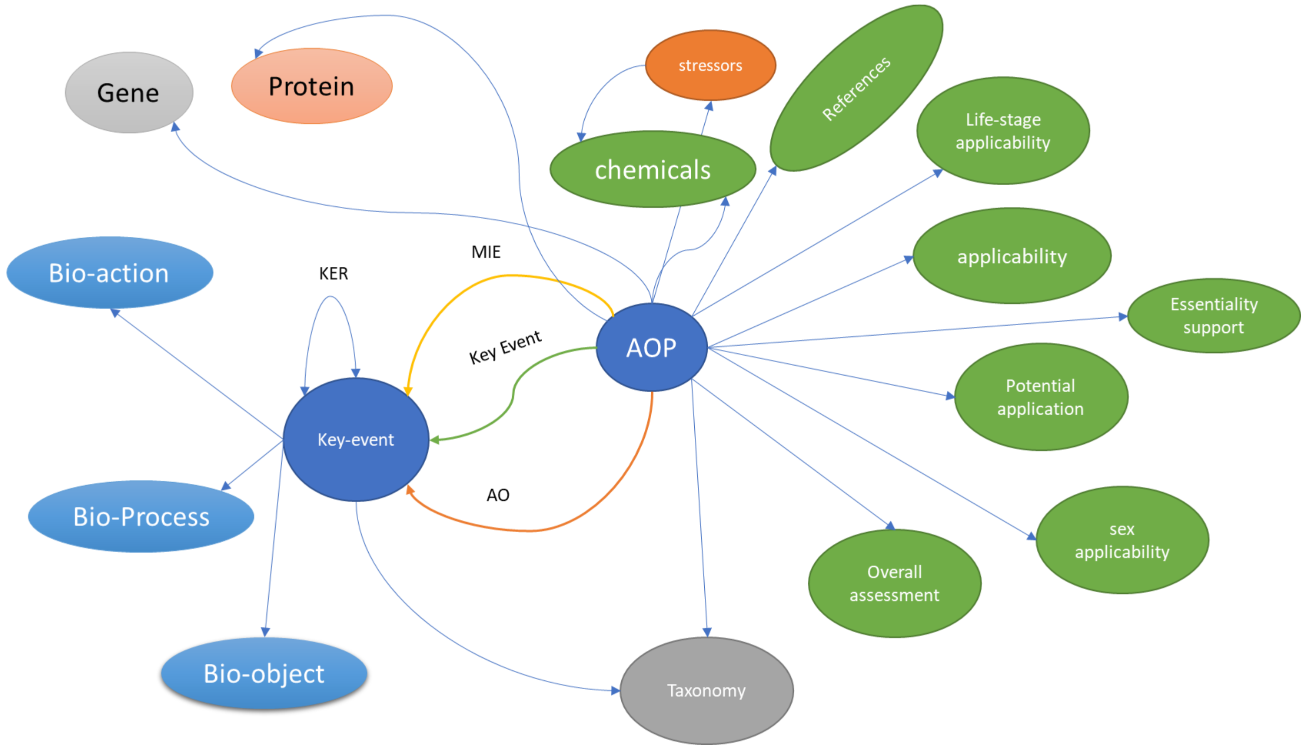


**Figure S2:** Adopted schema of AOP gives detailed information about the gene, chemical and disease mentioned in the textual description of AOPs.
